# Stress hyperglycemia exacerbates inflammatory brain injury after stroke

**DOI:** 10.1101/2024.05.14.594195

**Authors:** Seok Joon Won, Yiguan Zhang, Nicholas J. Butler, Kyungsoo Kim, Ebony Mocanu, Olive Tambou Nzoutchoum, Ramya Lakkaraju, Jacqueline Davis, Soumitra Ghosh, Raymond A. Swanson

**Author notes:** <u>Corresponding Author</u>: Raymond A. Swanson, (127) Neurology, SFVAMC, 4150 Clement St., Sn Francisco, CA 94121 USA.

## Abstract

Post-stroke hyperglycemia occurs in 30% - 60% of ischemic stroke patients as part of the systemic stress response, but neither clinical evidence nor pre-clinical studies indicate whether post-stroke hyperglycemia affects stroke outcome. Here we investigated this issue using a mouse model of permanent ischemia. Mice were maintained either normoglycemic or hyperglycemic during the interval of 17 - 48 hours after ischemia onset. Post-stroke hyperglycemia was found to increase infarct volume, blood-brain barrier disruption, and hemorrhage formation, and to impair motor recovery. Post-stroke hyperglycemia also increased superoxide formation by peri-infarct microglia/macrophages. In contrast, post-stroke hyperglycemia did not increase superoxide formation or exacerbate motor impairment in p47^phox-/-^ mice, which cannot form an active superoxide-producing NADPH oxidase-2 complex. These results suggest that hyperglycemia occurring hours-to-days after ischemia can increase oxidative stress in peri-infarct tissues by fueling NADPH oxidase activity in reactive microglia/macrophages, and by this mechanism contribute to worsened functional outcome.

---

Focal ischemic stroke results from occlusion of a cerebral artery or arteriole. Like other major injuries, stroke induces a systemic stress response. This stress response involves a sustained activation of the sympathetic nervous system, which in turn often causes elevated blood glucose (hyperglycemia) (1–4). The definition of hyperglycemia varies somewhat in clinical stroke studies, ranging from 6.1 to >10 mmol / L glucose, and by these criteria hyperglycemia is documented in 30% to 60% of patients following ischemic stroke (4–6). Some of these patients are diabetic, but most are not. Post-stroke hyperglycemia is strongly associated with negative outcome measures such as infarct size, mortality, disability, and poor recovery. This association is observed in ischemic stroke with or without thrombolysis, and it remains significant in patients with no prior history of hyperglycemia and in studies using logistic regression analysis to control for potentially confounding factors (6–9). However, clinical studies testing the effect of aggressive glucose control in patients after stroke found no improvement in functional outcome (4, 10), but an excess incidence of hypoglycemia in the treated groups leaves open the question as to whether post-stroke hyperglycemia is intrinsically deleterious. At present there are no established guidelines for management of stress hyperglycemia after stroke, despite its common occurrence (11).

Post-stroke hyperglycemia differs in several ways from hyperglycemia occurring during acute ischemia/reperfusion. The latter is unequivocally deleterious, as established by decades of both animal studies (12–15) and clinical observations (16–18). The mechanism by which hyperglycemia exacerbates acute ischemia/reperfusion injury stems in part from the role of glucose as an obligatory substrate for the production of the reactive oxygen species, superoxide and nitric oxide (19–24). The production of reactive oxygen species is particularly robust during acute reperfusion and re-introduction of oxygen and glucose into previously ischemic brain tissue, and the deleterious effects of hyperglycemia are reduced or absent in pre-clinical stroke models that do not involve reperfusion (25–27). Clinical data concur with this experimental observation (26, 28, 29). Thus, while the role of hyperglycemia in acute ischemia/reperfusion injury is well established, it remains uncertain if hyperglycemia has deleterious effects at later time points after ischemic stroke. It is even possible that hyperglycemia occurring at later time points might even be beneficial, e.g. by augmenting energy supply to marginally perfused brain tissue or suppressing spreading depression (30–33).

While reperfusion injury becomes progressively less likely to contribute injury with time after stroke, injury caused by the innate inflammatory response becomes a more important factor (34, 35). This immune response includes microglial activation, cytokine release, recruitment of circulating immune cells, and the release of proteases and reactive oxygen species. Proteases and reactive oxygen species in particular have been implicated in “bystander” death of neurons and other cell types after ischemic injury. Hyperglycemia exacerbates inflammation-induced injury in general (36), and in particular it promotes release of superoxide from activated immune cells (37, 38). Given the importance of inflammatory processes in secondary stroke injury (39), these observations suggest that hyperglycemia occurring many hours or days after stroke could significantly worsen outcomes after stroke. Surprisingly, the fundamental question as to whether post-stroke hyperglycemia has either beneficial or deleterious effects on stroke outcome has not previously been directly assessed in an experimental setting (40).

To address this gap, we evaluated the effects of hyperglycemia occurring hours-to-days after brain ischemia in non-diabetic mice. Our findings indicate that even a relatively short, 36-hour period of hyperglycemia that coincides with the peak brain inflammatory response after stroke significantly worsens both histological and functional outcomes, and that this deleterious effect is mediated at least in part by promoting microglial superoxide production.

## RESULTS

Mice subjected to photothrombotic cortical stroke developed an infarct in the targeted motor cortex (Fig. 1A). The infarct was accompanied by microglial activation in the peri-infarct tissue which, in agreement with prior reports (41), was robust by 24 - 48 hours after stroke (Fig. 1B). By 48 hours after stroke there was also a substantial increase in superoxide production in the peri-infarct tissue, which localized primarily to microglia / macrophages (Fig. 1 C,D).

**Figure 1.**
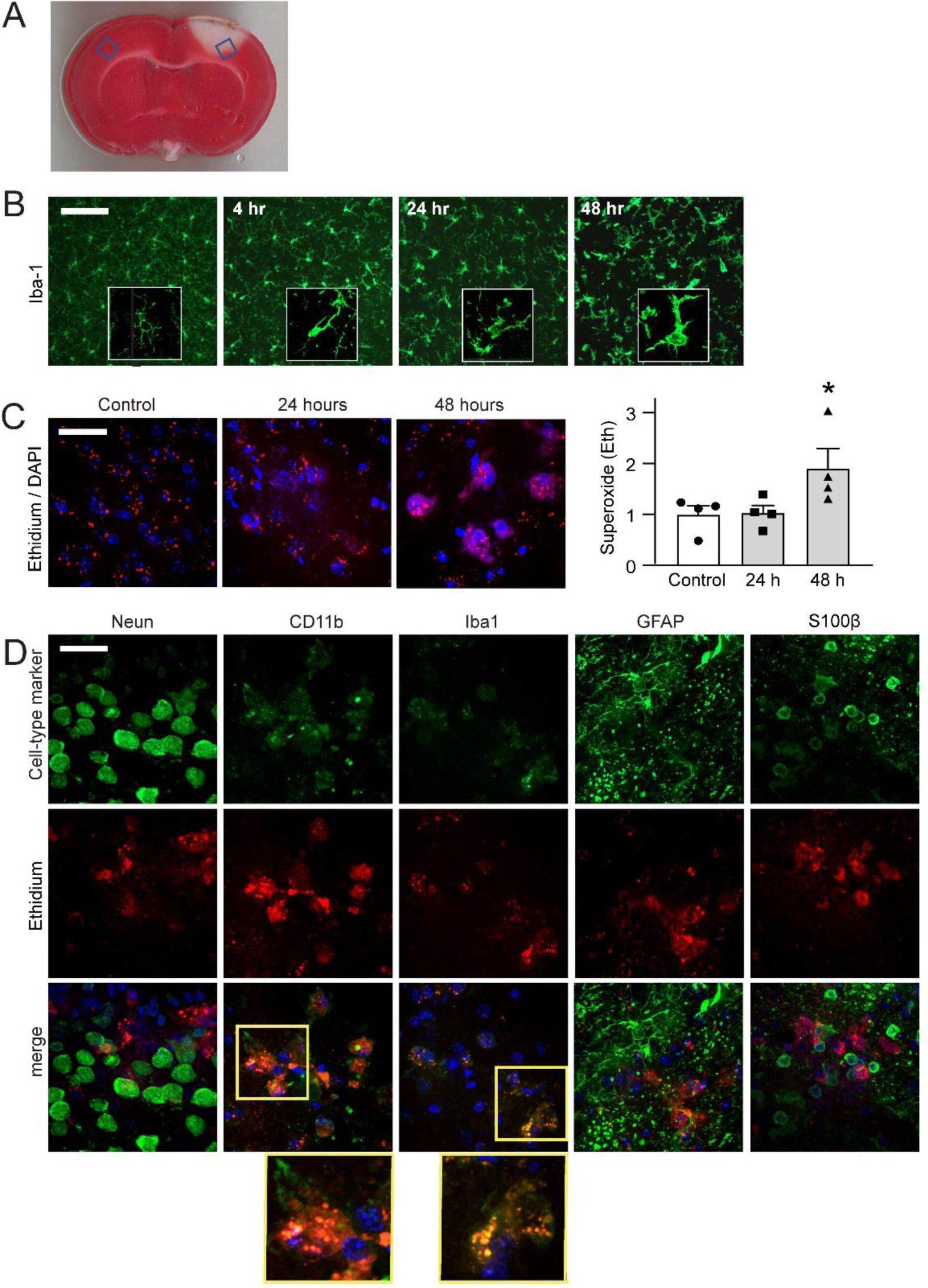
Delayed superoxide production by microglia/macrophages in peri-infarct cortex. (**A**) Representative TTC-stained coronal brain section through the area of infarct, with blue boxes showing the approximate areas photographed on each section. (**B**) Progressive change in peri-infarct microglial morphology over time after stroke. Microglia are immunostained for Iba-1 (green). (**C,D**) Increased superoxide production detected by oxidized ethidium (red) in the peri-infarct cortex at 48 hours after stroke. n = 4, *p < 0.05. Nuclei are counterstained with DAPI (blue). (**C**) Double labeling for oxidized ethidium fluorescence (Eth, red) and cell-type specific markers (green) for neurons (NeuN), microglia/macrophages (CD11b and Iba1) and astrocytes (GFAP and S100β). Cell nuclei are stained with DAPI (blue). Yellow boxes denote the areas magnified in adjacent panels. Images are representative of 4 mice under each condition. Scale bars = 20 µm.

Stroke-induced stress hyperglycemia usually persists for several days if untreated (4, 7). To determine if hyperglycemia at time points after the acute ischemic interval would increase the peri-infarct superoxide production, mice were given intraperitoneal glucose injections over a 3-hour interval beginning 45 hours after stroke. The glucose administration roughly doubled mean blood glucose levels (Fig. 2A,B), which corresponding to the magnitude of changes typically observed clinically in non-diabetic stress hyperglycemia (4, 7). The hyperglycemic mice showed a large increase in peri-infarct superoxide production relative to normoglycemic mice (Fig. 2 C,D). The increased superoxide production was accompanied by a corresponding increase in cellular oxidative stress, as assessed by the DNA damage marker γH2AX (Fig. 2 E,F). Double labeling with cell-type specific markers showed that the γH2AX foci were primarily in neurons (Supplemental Fig. S1).

**Figure 2.**
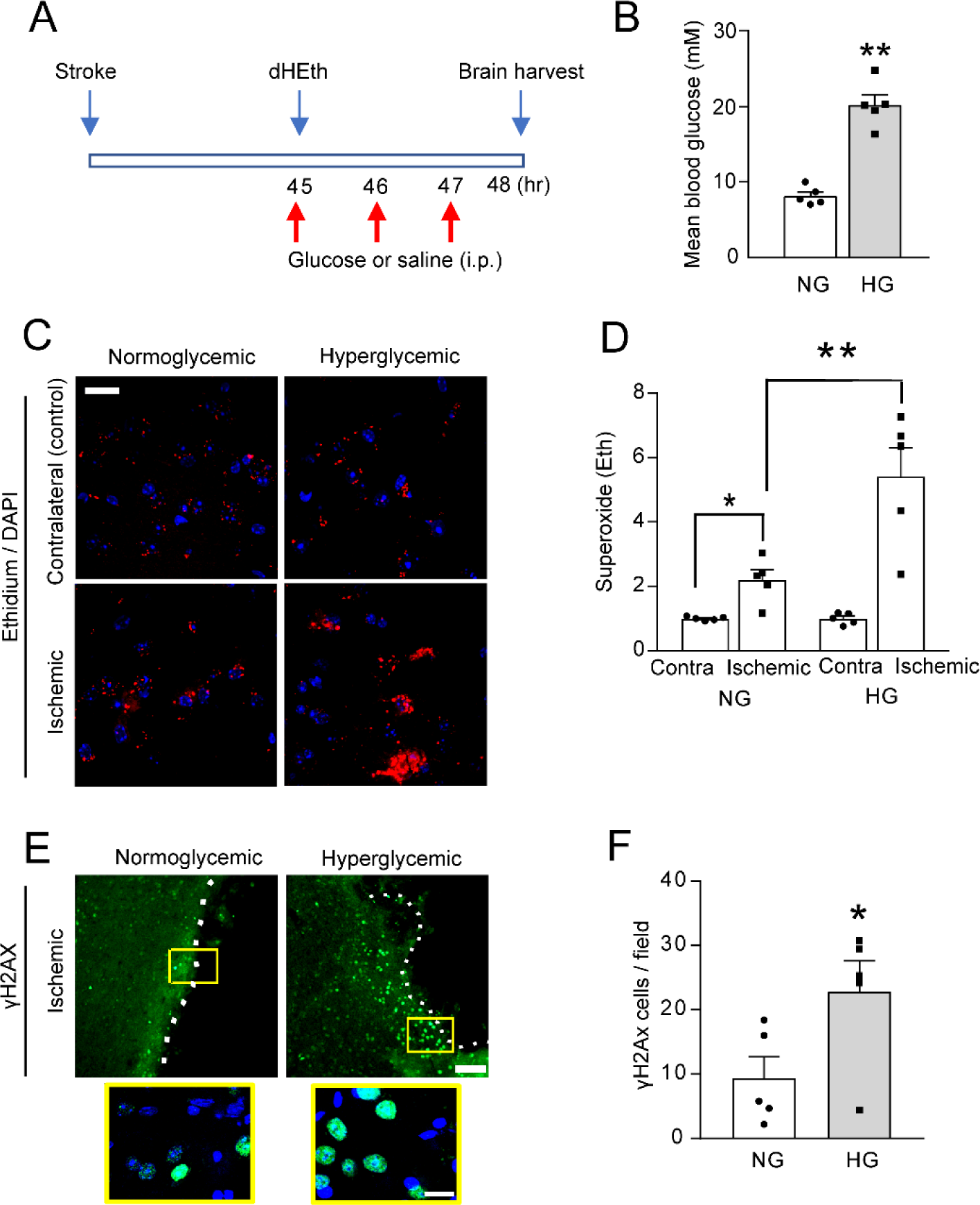
Hyperglycemia after stroke increases superoxide production in peri-infarct cortex. (**A**) Experimental design. Mice were injected i.p. with the superoxide indicator dihydroethidium (dHEth) and with glucose (or saline vehicle) beginning 45 hours after stroke, with brain harvest at 48 hours. (**B**) Mean blood glucose levels between 45 - 48 hours in normoglycemic (NG) and hyperglycemic (HG) mice. (**C,D**) Superoxide production as identified by oxidized ethidium fluorescence (red) in the peri-infarct and contralateral cortex of normoglycemic (NG) and hyperglycemic (HG) mice. Cell nuclei are stained blue. Scale bar = 20 µm. (**E,F**). DNA damage as evidenced by foci of γH2Ax formation (green) in the peri-infarct cortex of normoglycemic and hyperglycemic mice. Nuclei are stained blue. Lower panels are magnified views of the areas marked by white boxes. Scale bar = 100 µm (20 µm in the magnified images). Dotted lines denote infarct border. For all graphs, n = 5; **p < 0.01, * p < 0.05.

NADPH oxidase-2 is the primary source of superoxide produced by resident microglia and infiltrating macrophages (42). To confirm NADPH oxidase-2 as the source of superoxide in the peri-infarct brain, we treated mice with a peptide (gp91 ds-TAT) that inhibits assembly of the activated NADPH oxidase-2 complex (43). Mice injected with the gp91 ds-TAT peptide 4 hours prior to brain harvest showed a robust suppression of the superoxide signal, with no suppression of hyperglycemia (Fig. 3 A,B). This observation was corroborated in parallel study using p47^phox-/-^ mice, which are congenitally unable to form an active NADPH oxidase-2 complex. As expected, the p47^phox-/-^ mice likewise failed to show an increase in superoxide production in response to hyperglycemia (Fig. 3 C,D). Together, these results confirm NADPH oxidase-2 as the source of superoxide in the peri-infarct tissue and confirm the specificity of the dihydroethidium probe for superoxide in this experimental setting. The findings also comport with prior reports linking increased glucose availability to increased superoxide production via NADPH oxidase (19, 20, 44, 45).

**Fig. 3.**
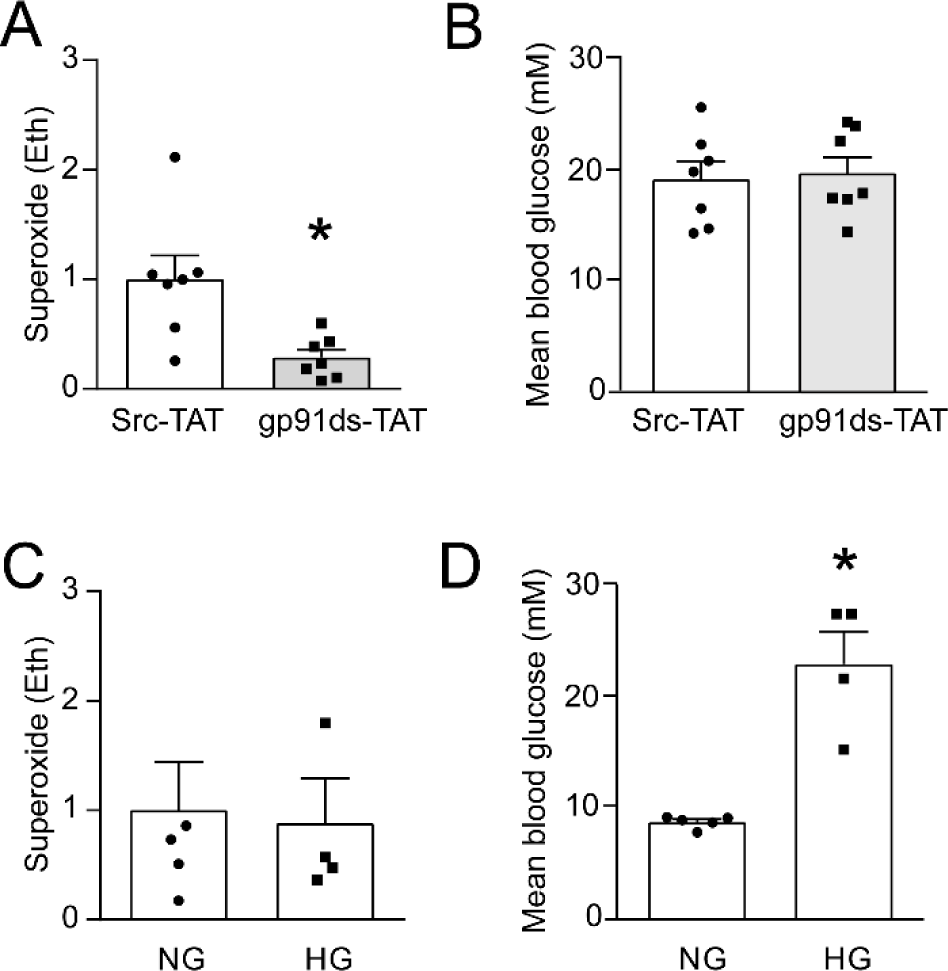
NADPH oxidase is the source of peri-infarct superoxide production. (**A,B)** The gp91ds-TAT peptide, which blocks NADPH oxidase-2 activity, prevents hyperglycemia-induced superoxide production without affecting blood glucose levels. **(C,D)** p47^phox-/-^ mice, which cannot form a functional NADPH oxidase-2 complex, show no increase in peri-infarct superoxide production when rendered hyperglycemic. n = 4-7, **p < 0.01. In A and C, oxidized ethidium fluorescence (Eth) measurements are normalized to the mean of the hyperglycemic group.

We next sought to determine whether hyperglycemia occurring in the post-stroke interval would affect histological or behavioral outcomes. Hyperglycemia in the clinical setting after stroke results from both reduced insulin secretion and accelerated hepatic glucose release (3). To mimic this effect in mice, we administered the α2-adrenergic agonist xylazine (46, 47) via osmotic pump, at a dose that had no discernible behavioral effect, in conjunction with glucose gavage feedings. This combination raised glucose levels to 2 - 3 times normal over the interval spanning 18 - 48 hours after stroke (Supplemental Fig. 2). At the 48-hour time point the osmotic pumps were removed and glucose gavage feeding was discontinued. Mice rendered hyperglycemic by this method were either euthanized at day 7 after stroke for histological assessments, or survived for an additional 3 weeks for behavioral assessments.

Histological assessments showed infarct volume to be modestly but significantly increased by post-stroke hyperglycemia (Fig. 4). There was also increased blood-brain barrier dysfunction in the peri-infarct tissue, of the mice rendered hyperglycemic, as assessed by IgG extravasation (Fig. 4 D,E), and this was accompanied by an increase in blood extravasation (Fig. 4 F,G). Of note, the hyperglycemic animals did not show elevated blood pressure relative to the normoglycemic mice (Supplemental Fig. 2), a factor that could independently promote post-stroke microhemorrhage.

**Figure 4.**
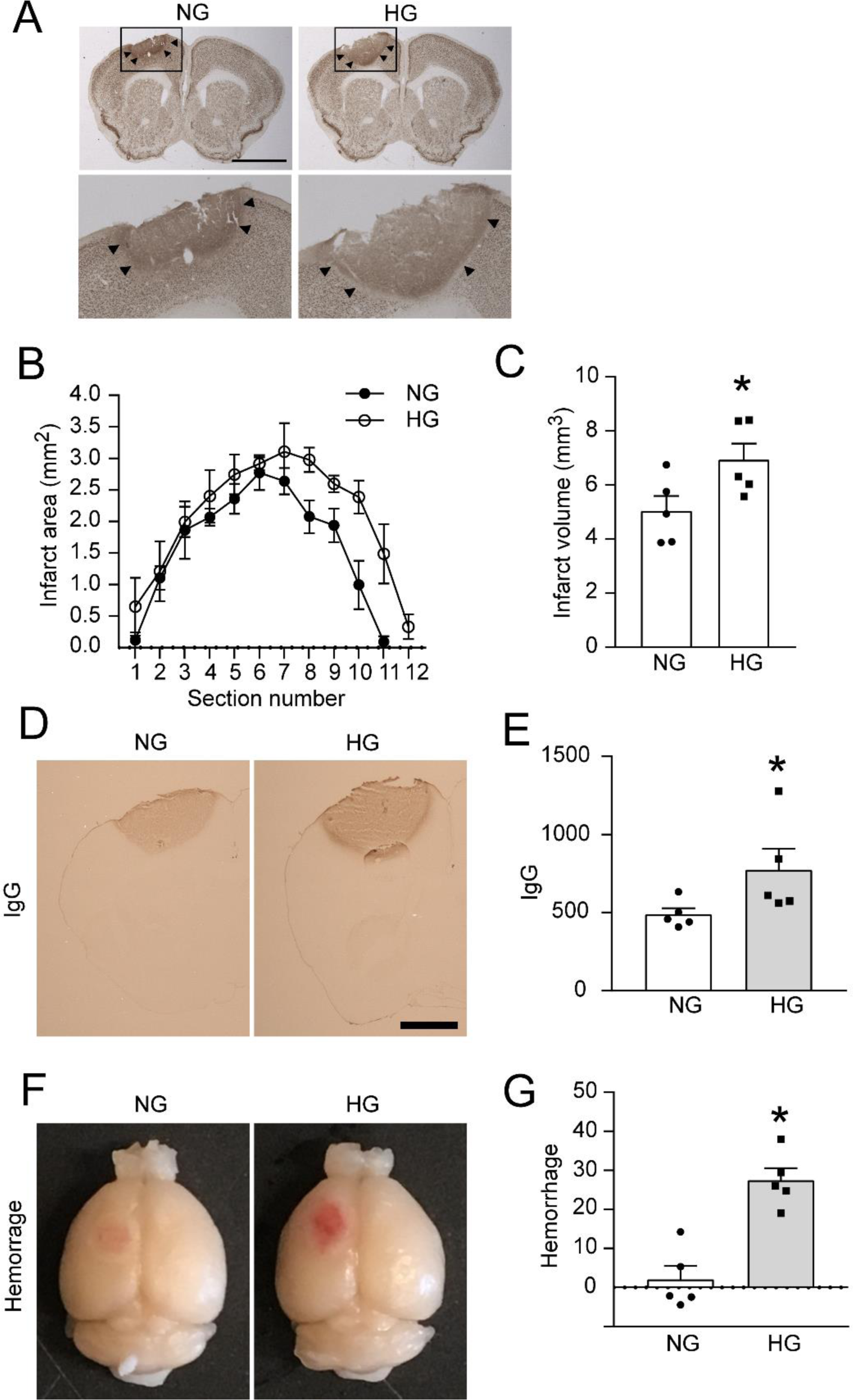
Delayed onset hyperglycemia after stroke increases infarct size, blood-brain barrier leakage, and cortical hemorrhage. (**A)** NeuN immunostaining in brains harvested 7 days after stroke. Areas marked with black boxes in low power view are shown in the magnified view below. (The infarcted tissue detached from the hyperglycemic (HG) but not normoglycemic (NG) brain sections.) Arrowheads indicate infarct edge. Scale bar = 2 mm. (**B, C**) Graphs show infarct areas in coronal sections and integrated infarct volumes in the normoglycemic and hyperglycemic treatment groups. (n = 5; \**p* < 0.05) (**D, E)** Blood-brain barrier leakage assessed by immunostaining for IgG. Scale bar = 1 mm. (n = 5; \**p* < 0.05). (**F,G)** Hemorrhage into infarct region in the two treatment groups. (n = 5; \*\**p* < 0.01, *p < 0.05).

Mice maintained for behavioral assessments were evaluated by the skilled reach test, sticky tape test, and rotating beam test over a 25 day period after stroke. In each of these studies, mice that had been rendered hyperglycemic between 18 - 48 hours after stroke recovered more slowly than the normoglycemic mice (Fig. 5 A-C). We then performed the behavioral studies in a cohort of p47^phox-/-^ mice, to test whether the worsened motor performance in the post-stroke hyperglycemic mice could be attributed to the increase in superoxide production caused by hyperglycemia. In contrast to results with the wild-type mice, the p47^phox-/-^ mice subjected to hyperglycemia did not show significantly impaired performance relative to the normoglycemic group, (Fig. 5 D-F).

**Figure 5.**
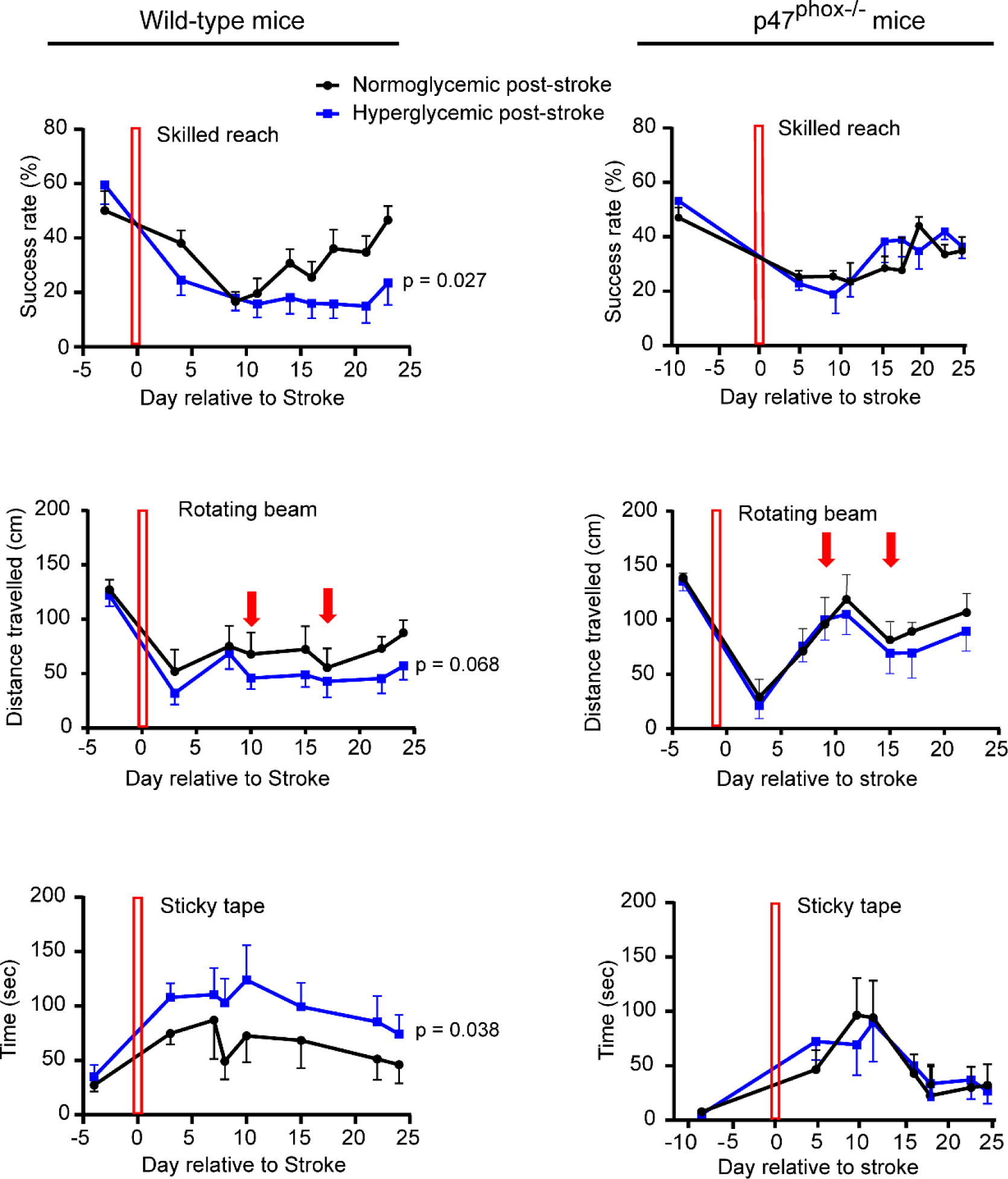
Delayed onset hyperglycemia after stroke impairs functional recovery in wild-type but not NADPH oxidase deficient mice. Red bars indicate day of stroke induction. Red arrows on rotating beam test indicate days that rotation rate was increased. * p < 0.05 values are based on repeated measures ANOVA comparing normoglycemic vs. hyperglycemia across all days post-stroke. n = 9 - 12 for wild-type mice, and n = 6 - 7 for p47^phox-/-^ mice.

The increase in superoxide production and oxidative stress induced after just 3 hours of hyperglycemia argues for a metabolic rather than transcriptional mechanism. However, hyperglycemia has also been shown to drive pro-inflammatory responses at the transcriptional level in activated microglia / macrophages (48–50). We therefore evaluated gene expression in peri-infarct cortex to determine if hyperglycemia might additionally act by a transcriptional mechanism to amplify the post-stroke inflammatory response. Mice were rendered hyperglycemic or normoglycemic in the 18 - 48 hour post-ischemic interval, and at the 48-hour time point brain tissue was taken from the peri-infarct cortex (or homologous region in sham stroke mice) and transcripts were evaluated for 767 genes (Supplementary Table 1). Genome-wide expression differences were observed between peri-infarct and control (sham stroke), as expected. By contrast, post-ischemic hyperglycemia produced relatively small changes (Fig. 6 A-C). To mitigate possible type II errors produced by multiple comparisons among hundreds of genes we pre-selected a small group of pro-inflammatory genes directly linked to cell injury for statistical analysis. Most of these showed large and statistically significant upregulation after stroke, but with none of these did hyperglycemia produce a significant further increase (Fig. 6 D). We also pre-selected 8 genes that putatively correlate with pro-inflammatory and anti-inflammatory microglial polarization (51), and here also found no statistically significant differences between the post-stroke hyperglycemic and post-stroke normoglycemic treatment groups (Fig. 6E). The full data sets of the gene expression studies results are provided in Supplemental Tables 1 - 3.

**Figure 6.**
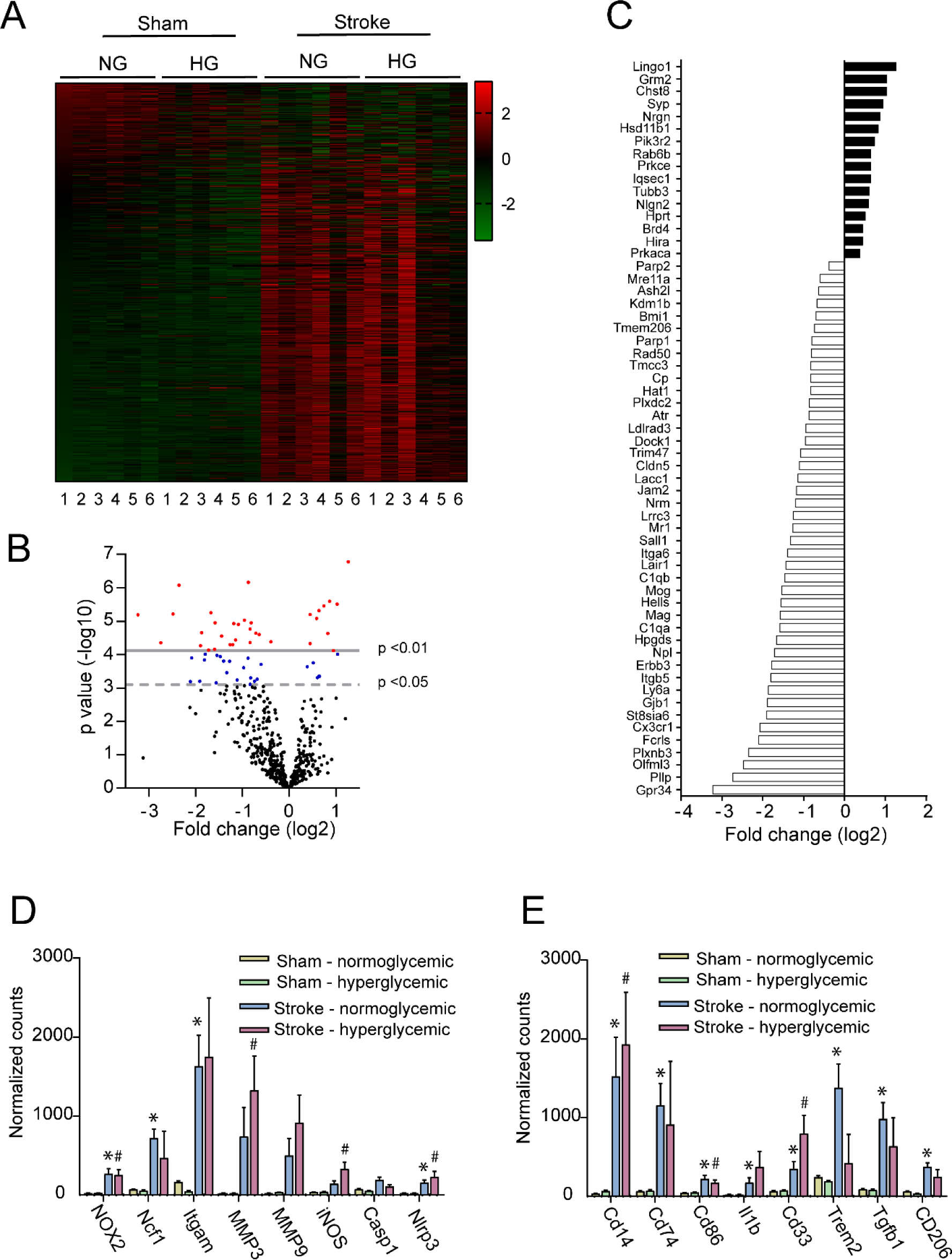
Effect of post-stroke hyperglycemia on gene expression in peri-infarct cortex. (**A**) Heatmap of gene expression in peri-infarct cortex. NG, normoglycemic; HG hyperglycemic. Vertical color bar indicates z-score calculated for each gene. Numbers below the heatmap refer to individual mice. (**B**) Volcano plot of heat map data showing effect of post-ischemic hyperglycemia on gene expression relative to the normoglycemic post-ischemic group. p values are not corrected for multiple comparisons. (**C**) Genes corresponding to those shown in (B) with at least p < 0.05 and > 1 fold change in expression. (**D**) Comparison of 8 pre-selected inflammation injury-associated genes among the 4 treatment groups. (**E**) Comparisons of selected genes associated with microglial pro-inflammatory and anti-inflammatory polarization. For (D) and (E), * p < 0.05 vs. sham stroke - normoglycemia and ^#^ p < 0.05 vs. sham stroke -, hyperglycemia after Bonferroni correction for multiple comparisons. There were no significant differences between the stroke - normoglycemic group and stoke - hyperglycemic group in either of the two pre-selected gene groups.

## DISCUSSION

Our studies found that hyperglycemia beginning many hours after stroke increased infarct size, blood-brain barrier disruption, hemorrhage formation, and motor dysfunction. These effects were accompanied by increased formation of superoxide by peri-infarct microglia / macrophages. In contrast, post-stroke hyperglycemia did not increase superoxide formation or exacerbate motor impairment in p47^phox-/-^ mice, which cannot form an active superoxide-producing NADPH oxidase-2 complex. Together, these findings indicate that hyperglycemia beginning many hours after stroke can increase oxidative stress in peri-infarct tissues by fueling microglial NADPH oxidase activity, and by this mechanism contribute to worsened functional outcome (Fig. 7).

**Figure 7.**
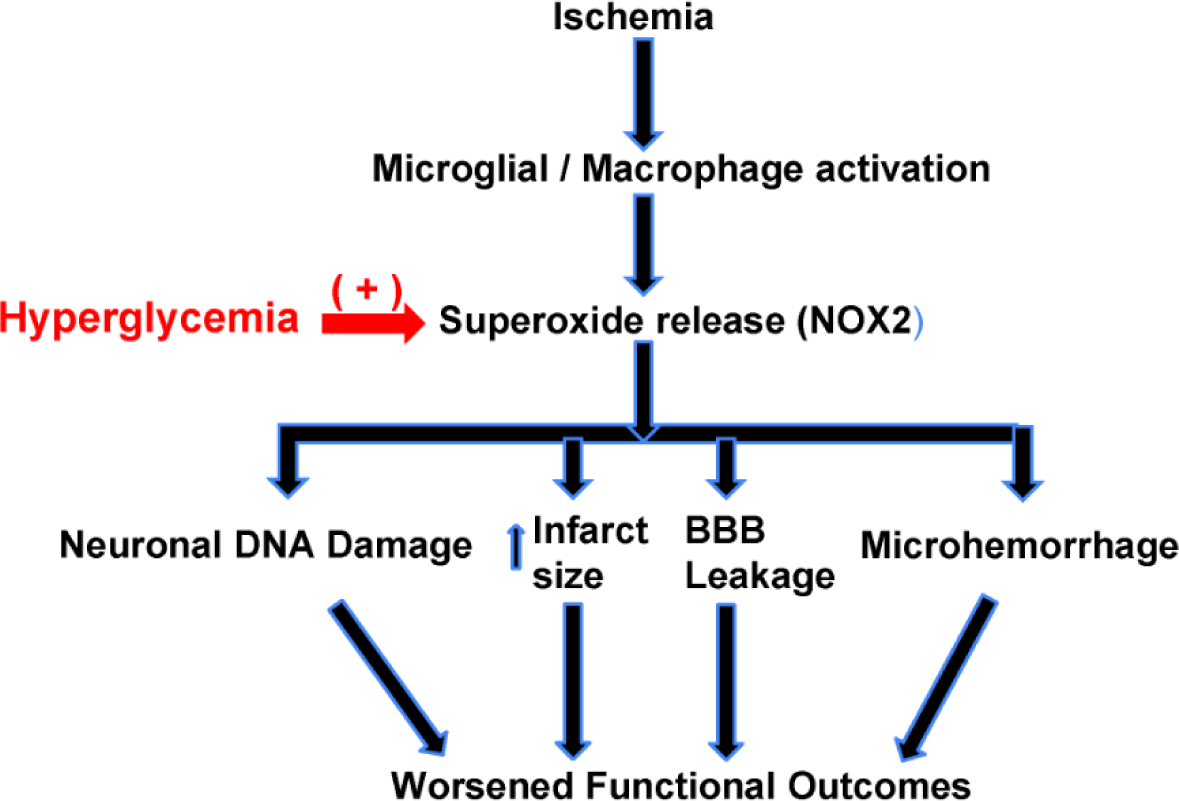
Relationships between brain inflammation, post-stroke hyperglycemia, and stroke outcomes. Hyperglycemia enhances provides glucose delivery to activated microglia / macrophages in peri-infarct brain. These cells utilize glucose for superoxide formation as part of the innate immune response to injury. Increased superoxide formation leads to increased oxidative stress in the peri-infarct tissue, which in turn leads to DNA damage, microvascular dysfunction, blood extravasation, infarct expansion, and impaired functional recovery.

The methods used to produce both stroke and hyperglycemia are important for the interpretation of the results obtained. We used photothrombotic permanent ischemia rather than an ischemia-reperfusion approach to produce stroke, thereby ensuring that effects of hyperglycemia could not be attributed to reperfusion injury. The photothrombotic approach also provided highly reproducible infarct size and location, which facilitates studies of motor impairment. The permanent ischemia model is also more germane to most clinical strokes, as the vast majority of these do not manifest either spontaneous or iatrogenic reperfusion in a time frame that can affect cell viability (52, 53). Prior studies indicate that the inflammatory response in the margins of a photothrombotic infarct are comparable to that observed in other stroke models (54).

It is difficult to maintain hyperglycemia for extended periods of time in non-diabetic experimental animals, a factor that likely contributes to the paucity of prior research in this area. In our initial set of studies, mice were rendered hyperglycemic on the day following stroke with repeated glucose administration. This served to prove principle that delayed hyperglycemia does increase oxidative stress in peri-infarct brain (and not elsewhere in brain), and that microglial NADPH oxidase was the primary source of the increased oxidative stress.

However, this approach cannot be used to produce a sustained glucose elevation, and in addition it provokes compensatory insulin release that may itself influence brain injury (55). Given that suppression of insulin release contributes to stress hyperglycemia, we instead adapted the approach of using low-dose xylazine administration - which acts at α2-adrenergic receptors to suppress insulin release - in combination with intermittent glucose administration. The levels of blood glucose obtained using both of these methods were roughly 2- 2.5 fold higher than mouse baseline, which is comparable to the magnitude of stress-induced hyperglycemia observed after stroke (4, 7).

A potential limitation to this method of inducing prolonged hyperglycemia is the possibility that xylazine could affect worsen stroke outcome by a mechanism other than its effect on blood glucose levels. The possibility of off-target effects cannot be entirely excluded, but increased superoxide production and oxidative stress were also observed in the study paradigm using glucose administration alone, in the absence of xylaxzine, and studies evaluating direct effects of α2-adrenergic agonists on brain injury have not observed injury exacerbation (56).

Superoxide release is a key aspect of the innate immune response. Though not intrinsically highly reactive, superoxide is converted in tissues to species such as peroxynitrite, hypochlorous acid, and hydroxyl radical, which are toxic not only to potential invading pathogens but also normal brain tissue. NADPH oxidase, specifically NADPH oxidase-2, is the primary source of superoxide generation by activated microglia and macrophages (42). Glucose is required for the production of superoxide, and also nitric oxide, because NADPH is the obligate substrate for the hexose monophosphate shunt that generates NADPH used as the electron donor for the enzymatic production of both of these reactive oxygen species and (21, 38). Elevated blood concentrations have been shown to increase superoxide production around ischemic brain, suggesting that delivery of glucose is be rate-limiting for superoxide production under ischemic conditions (23, 45). The oxidized ethidium signal used here to detect superoxide is not entirely specific for this reactive oxygen species, but the negation of this this signal in mice deficient in NADPH oxidase or treated with an NADPH oxidase inhibitor confirms that the ethidium production is attributable to superoxide or its downstream metabolites under the conditions of these studies.

Our tests of motor function showed impairment of contralateral forelimb after stroke with partial recovery over the following 3 weeks. All three motor tests employed showed impaired recovery in the mice rendered hyperglycemic after stroke, and crucially this effect of hyperglycemia was not observed in the p47^phox^ mice deficient in superoxide production. The mechanism by which post-stroke superoxide production impairs motor recovery is likely multifactorial. The observations presented here show increased neuronal DNA damage, blood-brain barrier dysfunction, microhemorrhage formation, and infarct size in the post-stroke hyperglycemic mice, each of which may be contributory factors.

Assessment of gene expression in peri-infarct cortex showed striking upregulation of several pro-inflammatory genes, but did not show statistically significant further elevations by hyperglycemia. This was unexpected, as prior reports have shown that glucose levels can influence pro-inflammatory gene expression in microglia / macrophages (48–50). It is possible that the signal from those cells was diluted by mRNA from other cell types, or that sampling at a different time point would have given a different result. Nevertheless, the lack of any observed additive effect of hyperglycemia on pro-inflammatory gene expression suggests that the primary mechanisms by which hyperglycemia influences peri-infarct superoxide production occurs at the metabolic rather than transcriptional level.

Together, results of this study demonstrate that hyperglycemia occurring many hours after stroke can negatively affect outcome through a mechanism involving superoxide production by per-infarct microglia / macrophages. The effects of hyperglycemia were observed under clinically relevant elevations in blood glucose (2 - 2.5 fold) and with relatively short hyperglycemia duration (30 hours). The findings also suggest that targeting peri-infarct superoxide production may be an effective intervention after stroke, particularly in patients exhibiting hyperglycemia.

## METHODS

### Animals

Wild-type C57BL/6 mice were obtained from the Jackson Laboratories. p47^phox-/-^ mouse were also obtained from the Jackson Laboratories, and subsequently back-crossed to wild-type C57BL/6 mice for >10 generations. Mice were used at age 3-5 months. A total of 109 male mice were used, 75 for histology studies and 34 for behavioral studies. Experimental group sizes were pre-determined to be n = 4 - 7 for histology studies and n = 6 - 12 for behavioral studies, based on the effect size and variances observed in prior similar studies. Six mice died during the behavioral testing phase of the study, all wild-type mice with post-stroke hyperglycemia, and data from these mice were excluded from analysis.

### Ischemic stroke

Mice were anesthetized with 2% isoflurane in 70% N_2_O/ balance O_2_, delivered through a ventilated nose cone. Rectal temperature was maintained at 37 ± 0.5° by using a homeothermic blanket throughout the surgical procedure. Photothrombotic permanent ischemia was induced by the Rose Bengal technique (57, 58). A PE10 polyethylene catheter (Becton Dickinson, Franklin, NJ) was introduced into the femoral vein, and the head was immobilized in a stereotaxic frame. The skull was exposed by skin incision, and a 2 mm diameter fiber optic cable was placed on the skull over the left primary motor cortex: 1.0 mm anterior, and 1.5 mm lateral to bregma. Rose Bengal (Sigma-Aldrich, St Louis, MO; 20 mg/kg) dissolved in saline was infused through the venous cannula for 1 minute, and vascular occlusion was induced by 10 minutes illumination with white light (KL 1500 LCD, SCHOTT North America Inc., Southbridge, MA) through the fiber optic cable. Light intensity and aperture diameter were set to produce an infarct that extended through cortex to the corpus callosum (Fig. 1A). Sham ischemia animals were injected with saline only but were otherwise treated identically. The incisions were sutured, bupivacaine (6 mg/kg) and buprenorphine (0.1 mg/kg) were administered subcutaneously, and the mouse was moved to a warm recovery chamber until awake and ambulatory.

### Induction of hyperglycemia

Mice were randomly assigned to hyperglycemia or normoglycemia treatment groups after the stroke surgery. Hyperglycemia was induced by two methods. For the initial studies, to achieve 3 hours of hyperglycemia, mice were administered three i.p injections of 40% glucose at one-hour intervals to produce blood glucose levels of 15 - 20 mM beginning 45 hours after stroke (Fig. 1B.) Normoglycemic mice received saline vehicle only. Blood glucose was detected in tail vein blood using an Accu-Chek Aviva Glucose meter at the indicated time points. To achieve more sustained hyperglycemia, mice were administered 10% glucose drinking water, access to a gel pack containing 3 ml of 30% glucose, and intermittent gavage feeding with 200 µl of 10 - 50% glucose beginning 17 hours after stroke (Supplemental Figure 2A). The concentration of glucose in the gavage was determined by the blood glucose concentration checked immediately beforehand at each time point: <13.8 mM, 50% glucose; 13.8 – 22.2 mM, 40% glucose; 22.2 – 27.7 mM, 30% glucose; >27.7 mM, 10% glucose. (Normal non-fasting blood glucose in the C57BL/6 mouse is approximately 9.4 mM (59).) In addition, the mice were given a continuous infusion of low-dose xylazine via subcutaneous osmotic mini-pump (Alzet model 2001; DURET Corporation, Cupertino, CA, USA) to suppress insulin secretion (60). The osmotic pumps were filled with xylazine (125 mg/ml) or distilled water vehicle and placed in the subcutaneous tissue of the upper back under isoflurane/ nitrous oxide anesthesia followed by subcutaneous bupivacaine (6 mg/kg) and buprenorphine (0.1 mg/kg) at termination of the surgery. Normoglycemic controls received standard drinking water and gel packs without glucose, were gavage fed with 200 ul of water only, and were implanted with osmotic pumps containing water only. The osmotic mini-pumps were removed under brief isoflurane/ nitrous oxide anesthesia at 31 hours after implantation (48 hours after stroke).

Where indicated, some mice were injected i.p. with 10 mg/kg gp91 ds-TAT 4 hours before the induction of hyperglycemia to inhibit the assembly of NADPH oxidase (43), or with a control TAT-conjugated peptide lacking any known biological effect.

### Blood pressure and blood glucose measurements

Blood pressures were measured in unanesthetized mice placed into a designated holder on a warming platform using a CODA noninvasive blood pressure measurement device (Kent Scientific Corporation, Torrington, CT) (61). Glucose was measured in 5 µl samples of tail vein blood using an Accu-Check blood glucose meter.

### Immunohistochemistry

Anesthetized mice were perfused with cold saline (0.9% NaCl) followed by phosphate-buffered 4% formaldehyde (PFA). After post-fixation with 4% PFA for 24 hours, brains were immersed for another 24 hours in 20% sucrose for cryoprotection. The brains were then frozen and 40 µm coronal sections were prepared with a cryostat. The fixed brain sections were pre-incubated in a blocking buffer (2% donkey serum and 0.1% bovine serum albumin in 0.1 M phosphate buffer) at room temperature for 30 minutes, and then incubated with the primary antibodies overnight at 4° C. The antibody sources and dilutions used are listed in Table 1. After washing, antibody binding was detected using fluorescent secondary antibodies listed in Table 1. Stained sections were mounted on glass slides in DAPI-containing mounting medium (Vector laboratories, Burlingame, CA). Control sections were prepared omitting either primary or secondary antibodies. In some cases antibody binding was visualized by the diaminobenzidine method (62) rather than with fluorescent secondary antibodies. For all histochemical quantifications, both the person taking the photographs and the person analyzing the images were blinded to the treatment condition.

**Table 1.** Primary antibody sources and dilutions used.

| Antibody | Source | ID | Host | Dilution |
| --- | --- | --- | --- | --- |
| NeuN | Millipore, Temecula, CA, USA | MAB377 | Mouse | 1:1000 |
| NeuN | Millipore, Temecula, CA, USA | ABN78 | Rabbit | 1:1000 |
| Iba1 | Abcam, Cambridge, MA, USA | Ab107159 | Goat | 1:1000 |
| CD11b | Bio-Rad, Hercules, CA, USA | MCA711 | Rat | 1:250 |
| GFAP | Millipore, Temecula, CA, USA | MAB360 | Mouse | 1:1000 |
| S100 | Abcam, Cambridge, MA, USA | Ab76729 | Rabbit | 1:100 |
| Mouse IgG | Vector Lab, Burlingame, CA, USA | AI-9200 | Mouse | 1:250 |
| anti-mouse IgG 405 | Thermo Fisher Scientific, Waltham, MA, USA | A31553 | Goat | 1:1000 |
| anti-mouse IgG 488 | Thermo Fisher Scientific, Waltham, MA, USA | A21202 | Donkey | 1:1000 |
| anti-rabbit IgG 488 | Thermo Fisher Scientific, Waltham, MA, USA | A21206 | Donkey | 1:1000 |
| anti-rat IgG 488 | Thermo Fisher Scientific, Waltham, MA, USA | A21470 | Rabbit | 1:1000 |
| anti-goat IgG 405 | Abcam, Cambridge, MA, USA | Ab175664 | Donkey | 1:1000 |
| anti-goat IgG 488 | Thermo Fisher Scientific, Waltham, MA, USA | A11055 | Donkey | 1:1000 |

### Superoxide detection

Dihydroethidium in 7.5% dimethyl sulfoxide / phosphate buffered saline pH 8.8 was administered i.p. at the indicated time points. After brain harvest and sectioning, 3 40 µm sections spaced 480 µm apart were analyzed from each animal through the infarct region. Six images spanning the infarct area were photographed with Zeiss fluorescence microscope (Observer. D1) at excitation 538 - 562 nm and emission 570 – 640 nm for detection of oxidized ethidium species (Eth) (63). In some cases, the sections were also immunostained for cell-type specific markers. Photographs were taken at predetermined regions of peri-infarct cortex, which was defined here as non-infarcted tissue within 250 µm of the lateral edge of the infarct as assessed by DAPI-stained nuclear morphology (Fig. 2A). The Eth fluorescence intensity signal was quantified using Image J software and expressed as a ratio of the integrated signal intensity in the peri-ischemic ischemic region relative to the homologous contralateral region, after subtraction of background signal. The background signal threshold was defined as the fluorescence intensity higher than 98% of pixels in the contralateral (non-ischemic) region, and pixels in both hemispheres that fell below this threshold were excluded from the analysis.

Results are presented as the average number of positive cells in each 168 µm x 168 µm photographed region from each animal.

### Quantification of γH2Ax-positive cells

Confocal images were taken at three predetermined peri-infarct locations on each of three sections from each animal. Cells were classified as positive if the mean γH2AX fluorescent signal was greater than that of 98% of pixels in the corresponding contralateral (non-ischemic) regions. Results are presented the average number of positive cells in each 168 µm x 168 µm photographed region from each animal.

### Quantification of IgG extravasation and hemorrhage

IgG extravasation was assessed in four brain sections spaced 480 µm apart through the infarct region (bregma +2.0 to +0.5). The sections were incubated with biotinylated anti-mouse IgG (H+L) for 1 hour at room temperature, followed by incubations with avidin-biotin complex, and diaminobenzidine (45). The integrated density of the IgG signal was measured over the entire lesioned hemisphere as area x (mean density - background mean density), with background mean density measured on the contralateral (non-injured) hemisphere. Results are presented as the average integrated density measured in the 4 sections from each animal.

Hemorrhage was quantified in photographs of the brain surface using Image J by defining 2 mm diameter regions of interest centered on the lesion site and the homologous contralateral site. The integrated densities were measured and expressed as a ratio of the lesioned site to the contra-lateral site for each brain.

### Infarct volume

Twelve brain sections spaced 240 µm apart, spanning the entire infarct region, were immunostained with anti-NeuN to identify infarcted tissue. The area of neuronal loss in each section was calculated in Image J software, and the infarct volume in each brain was calculated summing these for each animal and multiplying by the 2.88 mm span of the 12 sections.

### NanoString Gene expression analysis

Mice were perfused with saline and then brain tissues were collected from peri-infarct cortical tissue (2 mm thickness) surrounding the epicenter (2 mm diameter) of the lesion site using 2 mm and 4 mm punctures. Total RNA was extracted using High Pure RNA Tissue kit (Roche) following the manufacturer’s protocol. One hundred ng of RNA per sample were used for gene expression analysis using a NanoString nCounter (NanoString Technologies). The nCounter mouse neuroinflammation panel of 757 genes (Supplemental Table 1) was supplemented by 10 additional genes (NOX1, NOX2, NOX4, iNOS, MMP3, MMP9, IL-6, CCL22, CD206). Quality of the hybridized signals was evaluated by assessing binding density, imaging, and positive controls as recommended by the manufacturer. Data that passed these quality control steps was processed with background correction and normalized to synthetic positive control targets and a set of housekeeping genes across the entire data set. Statistical analysis and graph preparation was performed with the nSolver 4.0 software.

### Behavioral studies

Mice were divided into five cohorts: three cohorts of wild-type mice and two cohorts of p47^phox-/-^ mice, each containing 5-7 animals. Each cohort was evaluated with three tests of motor dexterity: the skilled reaching test, sticky tape test, and rotating beam test. The mice were acclimated to handling over a 2 - week period prior to stroke, during which time they were also acclimated to each of the behavioral tests. Post-stroke testing was initiated 3 days after stroke and repeated twice-weekly for 3 weeks.

For the Wishaw skilled reaching task we used custom-made automated box as previously described for rats (64, 65), size-modified for use in mice. The mice were trained to reach through a 0.5 cm slit for a 14 mg rodent diet pellet (Bio-Serv, # F0071) prior to induction of stroke. Paw preference was noted during initial training sessions, and reaching slit was placed to facilitate use of the preferred paw. Mice were fasted overnight prior to the training sessions and fed 10% of their body weight after the training (minus the amount eaten during training). Mice were considered adequately trained when they were able to successfully grab and eat a pellet in 30 of 60 consecutive trials on two consecutive training sessions. Strokes were induced the hemisphere contralateral to the dominant paw. During post-stroke testing mice were presented with 60 opportunities to grab a pellet, and the percent of successful grabs (pellets eaten) was scored. A trial was not scored if the mouse failed to make at least 30 attempts.

The rotating beam test (66) requires mice to traverse the length of a rotating 145 cm long / 1 cm diameter beam. The task is scored by the distance the mouse covers before falling off onto a safety net below. The mice were habituated to the rotating beam for 4 days, with rotation speed increased from 0 rpm to 9 rpm. The mice completed 4 repetitions on the beam on each day of training: 2 clockwise and 2 counter-clockwise. The beam was wiped clean with bleach between mice. Mice were considered adequately habituated when able to traverse the full 145 cm at a 9 rpm rotation speed on 3 consecutive trials. Post-stroke evaluations were begun on day 3 after stroke and repeated twice-weekly for 3 weeks. The beam rotation speed was set at 9 rpm on days 1 - 10 after stroke and then increased as the animals improved performance; to 15 rpm and then to 20 rpm. Each testing session consisted of 4 repetitions on the beam: 2 clockwise and 2 counter-clockwise. The mean distance traveled was recorded as the score obtained for each testing day.

The sticky tape test was initiated by placing each mouse in a beaker for 1 minute to acclimate. A small sticker was then placed on the paw contralateral to the stroke. Time required to completely remove the sticker was recorded. Mice that had not removed the sticker within 120 seconds were assigned a time of 120 seconds. Trials were run twice each testing day with at least 15 minutes in between each trial, and the average of the two trials was recorded. The beaker was washed with 70% EtOH in between mice.

### Statistics

All data are expressed as means ± s.e.m., with the “n” of each experiment defined as the number of mice or, for cell cultures, the number of independent experiments. Behavioral outcome data were assessed by repeated measures ANOVA. The pre-selected gene expression comparisons were evaluated by ANOVA with Bonferroni correction, and the P values for all other genomics data were evaluated by the Benjamini-Yekutieli method (67). All other data were analyzed using two-sided T-tests.

### Study approval

Studies were approved by the animal studies committees at the San Francisco Veterans Affairs Medical Center, and were performed in accordance with the National Institutes of Health Guide for the Care and Use of Laboratory Animals and the ARRIVE guidelines.

### Data Availability

All primary genomic data is included in appendices to this publication. Primary data pertaining to the histology and behavioral studies will be provided upon email request to the corresponding author.

## AUTHOR CONTRIBUTIONS

SJW designed studies, performed experiments, analyzed data, and drafted the manuscript; YZ performed experiments; NJB performed experiments and analyzed data; KK designed and produced the mouse skilled reaching box; EM performed experiments; OTN performed experiments; RL performed experiments; JD performed experiments and analyzed data; SG performed experiments; RAS supervised the project, analyzed data, and edited the final manuscript.

## ACKNOWLEDGEMENTS

This work was supported by the NIH (R01 NS081149, RAS), and by the Dept. of Veterans Affairs. These sponsors were not involved in study design, data analysis, decision to publish, or other aspects of this work. We thank Todd C. Peterson, Dept. Psychology, University of North Carolina for advice on the rotating beam behavioral assessment, and Rebecca Fong for expert technical assistance.

