## Supplemental Figures for "Stress hyperglycemia exacerbates inflammatory brain injury after stroke"

### SUPPLEMENTARY FIGURES

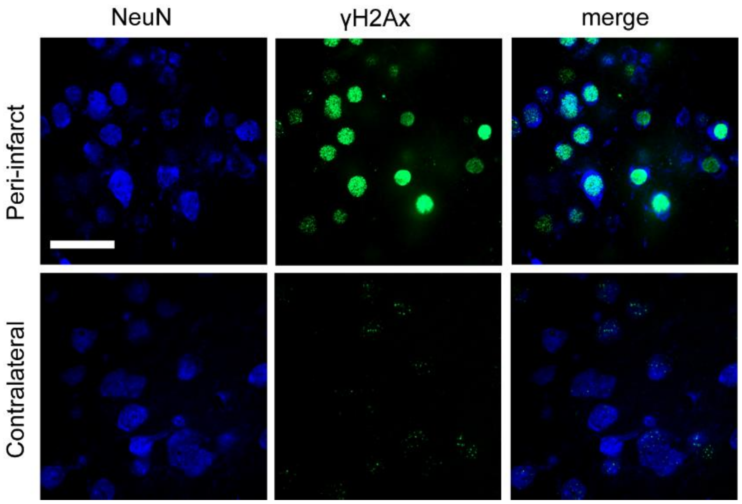


**Supplemental Figure S1. DNA damage in peri-infarct neurons.**

Images are from peri-infarct cortex of mice rendered hyperglycemic as in Fig. 1. Neuronal nuclei (as labeled by NeuN, blue) show co-localization with the DNA damage marker γH2AX (green), whereas neuronal nuclei in the contralateral, non-injured cortex do not. Representative of images obtained from 4 mice. Scale bar = 20 µm.

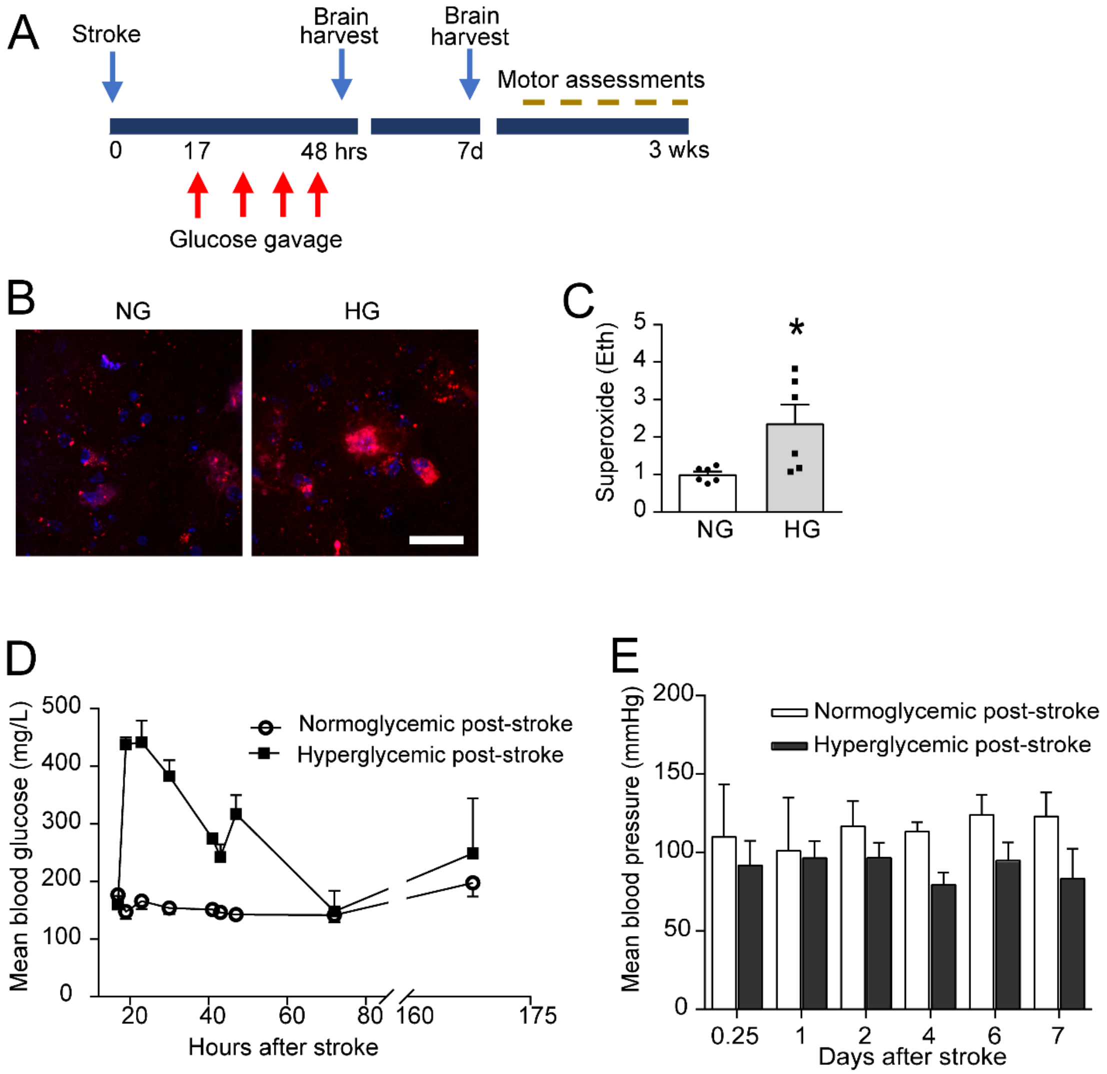


**Supplemental Figure S2. Experimental design for long-term end point studies.**

(**A**) Mice received glucose by 4 gavage feedings between 17 and 48 hours after stroke, along with xylazine via osmotic pump to attenuate insulin secretion. One cohort of mice was euthanized at either 48 hours or 7 days for brain histology, and a second cohort was evaluated over 3 weeks with assessments of motor function. (**B, C**) Mice rendered hyperglycemic showed increased superoxide production at in the peri-infarct cortex at the 48-hour time point. Images show Eth fluorescence (red), with cell nuclei stained blue. Scale bar = 20 µm. (n = 6; $*p < 0.05$) (**D**) Mean blood glucose values in the hyperglycemic and normoglycemic mice (n = 4). (**E**) Post stroke mean arterial blood pressures in the normoglycemic and hyperglycemic mice (n = 4).

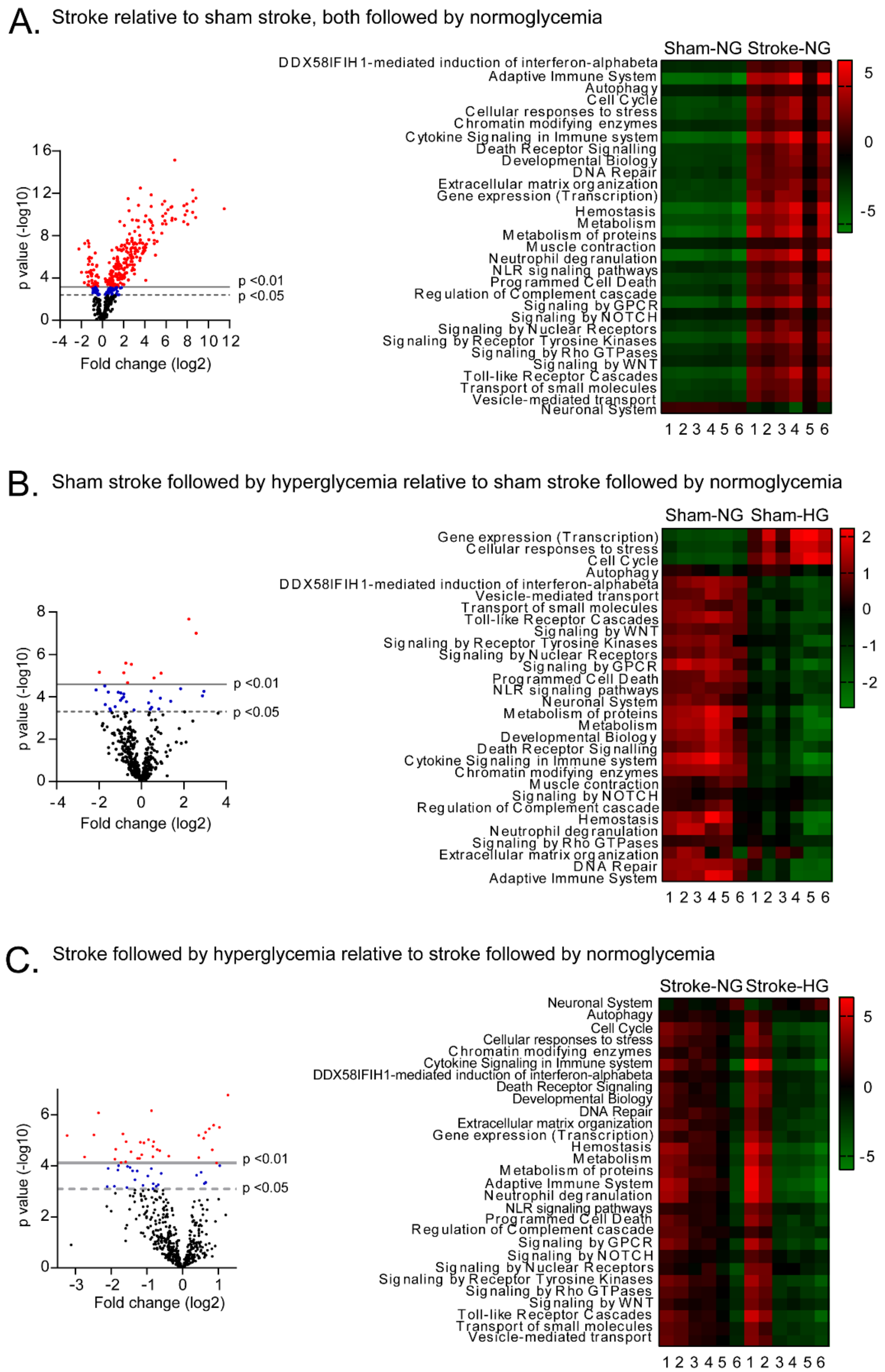


**Supplemental Figure S3**. **Pathway analysis of stroke and hyperglycemia effects on gene expression in brain cortex.**

**(A**) Volcano plot of gene expression changes induced by stroke relative to sham stroke, both followed by normoglycemia. Heatmap shows pathway analysis of this comparison. (**B**) Volcano plot of gene expression changes induced by hyperglycemia relative to normoglycemia after sham stroke. Heatmap shows pathway analysis of this comparison. (**C**) Heatmap shows pathway analysis of gene expression changes induced by stroke followed by hyperglycemia relative to stroke followed by normoglycemia. The volcano plot of these gene expression changes is shown in figure 6B. Numbers below each heatmap designate individual mice, and vertical color bars indicate Z-score. Tabulated data for panels A-C are provided in Supplementary Table 3.
